# Adipokinetic hormone signaling mediates the enhanced fecundity of *Diaphorina citri* infected by ‘*Candidatus* Liberibacter asiaticus’

**DOI:** 10.1101/2023.10.24.563719

**Authors:** Jiayun Li, Paul Holford, George Andrew Charles Beattie, Shujie Wu, Jielan He, Shijian Tan, Desen Wang, Yurong He, Yijing Cen, Xiaoge Nian

**Affiliations:** National Key Laboratory of Green Pesticide, Department of Entomology, College of Plant Protection, South China Agricultural University, Guangzhou, 510642, China; Henry Fok School of Biology and Agriculture, Shaoguan University, Shaoguan, 512005, China; School of Science, Western Sydney University, Locked Bag 1797, Penrith, NSW 2751, Australia

**Keywords:** huanglongbing, *Diaphorina citri*, *DcAKH*/*DcAKHR*, miR-34, mutualism

## Abstract

*Diaphorina citri* serves as the primary vector for ‘*Candidatus* Liberibacter asiaticus’ (*C*Las), the bacterium associated with the severe Asian form of huanglongbing. *C*Las-positive *D. citri* are more fecund than their *C*Las-negative counterparts and require extra energy expenditure. Therefore, understanding the molecular mechanisms linking metabolism and reproduction is of particular importance. In this study, we found adipokinetic hormone (*DcAKH*) and its receptor (*DcAKHR*) were essential for increasing lipid metabolism and fecundity in response to *C*Las infection in *D. citri.* Knockdown of *DcAKH* and *DcAKHR* not only resulted in the accumulation of triacylglycerol and a decline of glycogen, but also significantly decreased fecundity and *C*Las titer in ovaries. Combined *in vivo* and *in vitro* experiments showed that miR-34 suppresses *DcAKHR* expression by binding to its 3’ untranslated region, whilst overexpression of miR-34 resulted in a decline of *DcAKHR* expression and *C*Las titer in ovaries and caused defects that mimicked *DcAKHR* knockdown phenotypes. Additionally, knockdown of *DcAKH* and *DcAKHR* significantly reduced juvenile hormone (JH) titer and JH signaling pathway genes in fat bodies and ovaries, including the JH receptor, *methoprene-tolerant* (*DcMet*), and the transcription factor, *Krüppel homolog 1 (DcKr-h1)*, that acts downstream of it, as well as the egg development related genes *vitellogenin 1-like* (*DcVg-1-like*), *vitellogenin A1-like* (*DcVg-A1-like*) and the vitellogenin receptor (*DcVgR*). As a result, *C*Las hijacks AKH/AKHR-miR-34-JH signaling to improve *D. citri* lipid metabolism and fecundity, while simultaneously increasing the replication of *C*Las, suggesting a mutualistic interaction between *C*Las and *D. citri* ovaries.

## Introduction

The majority of devastating plant pathogens, including viruses and bacteria, are transmitted by insect vectors [1]. In recent decades, the interactions between vectors and pathogens have been widely studied [1-4]. However, studies on the mechanisms underlying these interactions have mainly focused on vector-virus or vector-fungus interactions; less research has been performed on vector-bacteria interactions. Knowledge of the relationships between vectors and pathogens is of great significance for understanding the epidemiology of plant pathogens, and this knowledge may help develop new strategies for controlling both vectors and pathogens. The severe Asian form of huanglongbing (HLB) is currently the most destructive citrus disease in Asia and the Americas where it is associated with the phloem-restricted, Gram-negative, α-Proteobacteria ‘*Candidatus* Liberibacter asiaticus’ (*C*Las) and transmitted by the Asian citrus psyllid, *Diaphorina citri* Kuwayama (Hemiptera: Psyllidae) [5-9]. However, since *C*Las has not been cultured *in vitro,* Koch’s postulates have not been proven. Control of *D. citri* is the most effective way to prevent HLB epidemics [6,7] but experience in Asia [10] and the United States of America [11,12], and recent press reports from Brazil, indicate that it is impossible to suppress psyllid populations with insecticides to levels that prevent spread of *C*Las.

Previous studies have reported that infection with *C*Las significantly increases the fecundity of *D. citri* [13-15]. Reproduction is a costly process in terms of energy usage in the adult life of female insects [16], and there is an inevitable trade-off between lipid storage and use, because animals mobilize their lipid reserves during reproduction. However, the mechanism of how *D. citri* maintains a balance between lipid metabolism and increased fecundity after infection with *C*Las remains unknown. In insects, energy mobilization is under control of adipokinetic hormone (AKH), which was the first insect neurohormone to be identified; it was shown to stimulate lipid mobilization for locomotor activity in locusts [17]. To date, more than 80 different forms of AKH have been identified or predicted in arthropod species due to similar structural characteristics of the enzyme [18]. The mechanisms of synthesis and biological activity of AKH have also been investigated. Insect store lipids in the form of triacylglycerides (TAG) and as carbohydrates in the form of glycogen in their fat bodies [19]. Under conditions of energy demand, AKHs are synthesized in the corpora cardiaca, released into the hemolymph and bind to receptors (AKHRs) located on the plasma membrane of fat body cells. AKHR binding and activation triggers the release of energy-rich substrates after which various cellular signaling pathway components come into play [20]. AKHRs are a class of G protein-coupled receptors and were first identified in *Drosophila melanogaster* [21] and *Bombyx mori* [22]; subsequently, they were identified in a large number of insect species [23-26].

While the AKH pathway is recognized for its role in mobilizing stored lipids, its involvement in vector-pathogen interactions remains understudied. For example, in *Anopheles gambiae*-*Plasmodium falciparum* interactions. *P. falciparum* infection influences *A. gambiae* AKH signaling, lipid metabolism, and mobilization of energy reserves to support the parasite’s metabolic and structural needs [27]. In another example, infection of cockroaches with the fungus *Isaria fumosorosea* result in a significant rise in AKH levels in the central nervous system in response to stress responses [28]. The functions of the AKH/AKHR in insect-pathogen interactions need further investigation. Recent studies have shown that AKH/AKHR signaling plays a regulatory role during female reproduction in insects. For example, depletion of AKHR in *Bactrocera dorsalis* resulted in TAG accumulation, decreased sexual courtship activity, and fecundity [29]. AKHR knockdown in *Nilaparvata lugens* interferes with trehalose homeostasis and vitellogenin uptake by oocytes, which causes delayed oocyte development and reduced fecundity [30]. AKH/AKHR signaling also regulates vitellogenesis and egg development in *Locusta migratoria* via triacylglycerol mobilization and trehalose homeostasis [31]. As outlined above, the fecundity of *D. citri* is increased by *C*Las. Whether AKH/AKHR signaling participates in this increase in fecundity due to infection via alterations in lipid metabolism is not currently known.

Although numerous studies have been conducted on the AKH/AKHR signaling pathway, miRNA-mediated regulation of the AKH signaling pathway at the post-transcriptional level has been little studied. MicroRNAs (miRNAs) are small, non-coding RNAs containing approximately 22 nucleotides that act mostly on the 3’ untranslated region (UTR) of target mRNAs, resulting in translation repression or mRNA degradation [32]. miRNAs play important roles in regulating cellular events at the post-transcriptional level during biological processes [32,33,34]. For example, in *D. citri*-*C*Las interaction, *C*Las hijacks the JH signaling pathway and host miR-275 that targets the *vitellogenin receptor* (*DcVgR*) to improve *D. citri* fecundity, while simultaneously increasing the replication of *C*Las itself, suggesting a mutualistic interaction in *D. citri* ovaries with *C*Las [35]. In term of host-virus interactions, miR-8 and miR-429 target *Broad isoform Z2*, the gene involved in the hormonal regulation of single nucleopolyhedrovirus (HaSNPV)-mediated climbing behavior in *Helicoverpa armigera* [36]. In relation to reproduction, in *Aedes aegypti*, miR-275 is essential for egg development [37], miR-309 for ovarian development [38] and miR-8 for other reproductive processes [39]. As for lipid metabolism, knockdown of miR-277 in *A. aegypti* activates insulin/FOXO signaling by increasing the expression of ilp7 and ilp8, which reduces TAG levels and inhibits ovarian development [40]. Nevertheless, the functions of the miRNAs in insect-pathogen interactions need further investigation. To date, there are no reports that miRNAs participate in *D. citri*-*C*Las interactions or are hijacked by *C*Las to affect lipid metabolism and the fecundity of *D. citri*. As a consequence, in this study, we used the *D. citri*-*C*Las system as a model to study the molecular events associated with AKH/AKHR signaling that affect lipid metabolism and increase fecundity of *D. citri* induced by *C*Las. Our study contributes to our understanding of the interactions between vectors and pathogens and may also provide new ways for controlling *D. citri* and HLB.

## Results

### Effects of *C*Las infection on the reproduction and metabolism of adult *D. citri*

Total TAG and glycogen levels in fat bodies of *C*Las-negative and *C*Las-positive psyllids were determined. After infection with *C*Las, the TAG and glycogen levels of *C*Las-positive psyllids significantly increased compared with *C*Las-negative individuals (Figures 1A-B). Since TAG is mainly stored in fat bodies as lipid droplets, we next evaluated the changes in the lipid droplet size using Nile red staining. As shown in Figure 1C, the size of lipid droplets in the *C*Las-positive group was significantly larger than those of the *C*Las-negative group. In order to evaluate the effects of *C*Las on the reproduction of *D. citri*, the preoviposition period, oviposition period and fecundity of both *C*Las-negative and *C*Las-positive psyllids were examined. After infection with *C*Las, the preoviposition period was significantly shortened, the oviposition period was extended, and the fecundity was markedly increased (Figures 1D-E).

**Figure 1.**
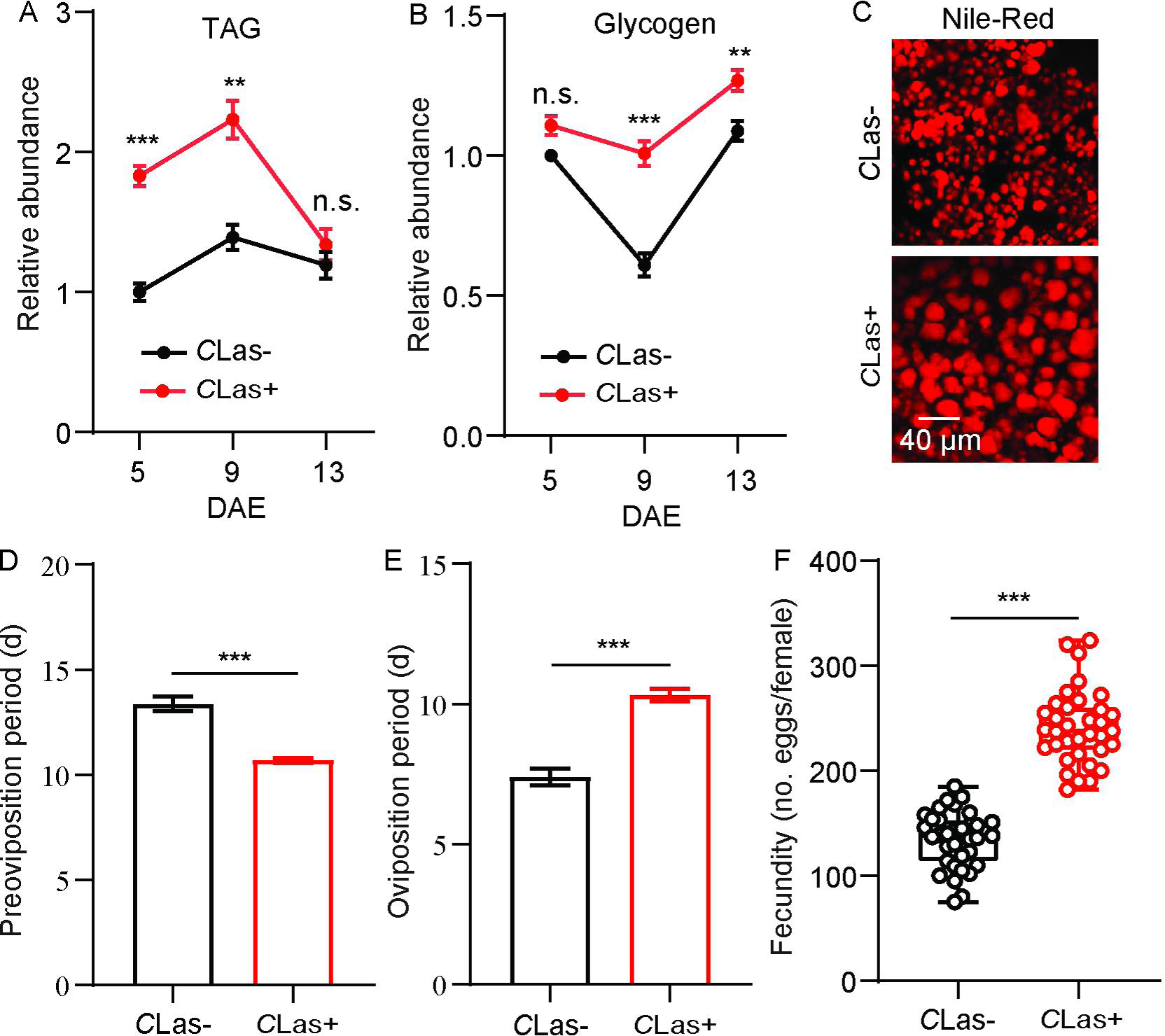
Effects of *C*Las on lipid metabolism and reproductive behavior of *D. citri*. (A) Comparison of TAG levels in the fat bodies of *C*Las-positive (*C*Las+) and *C*Las-negative (*C*Las-) females 5, 9, 13 days after emergence (DAE). (B) Comparison of glycogen levels in the fat bodies of *C*Las+ and *C*Las-females 5, 9, 13 DAE. (C) Lipid droplets in fat bodies dissected from *C*Las+ and *C*Las-females 9 DAE stained with Nile red. Scale bar = 40 μm. (D-F) Comparison of the preoviposition period, oviposition period and the fecundity of *C*Las+ and *C*Las-adults. Data are shown as means ± SEM. The significant differences between *C*Las-positive and *C*Las-negative psyllids were determined using Student’s *t*-tests (\*\**p* < 0.01, \*\*\**p* < 0.001).

### *DcAKH* is involved in changes in metabolism and fecundity in *D. citri* mediated by *C*Las

Using our transcriptome database, cDNA sequences of *DcAKH* were identified and cloned. The cDNA sequence of *DcAKH* is 225 bp in length, encodes a deduced polypeptide of 74 amino acid residues and has the conserved structural characteristics of a neuropeptide. The conserved structure was “signal peptide + mature peptide + related peptide”, and the mature peptide sequence of *Dc*AKH is “QVNFSPNW” (Figure S1A). A phylogenetic analysis was conducted to evaluate the association of *Dc*AKH with other insect AKHs; *Dc*AKH was most closely related to the AKHs of three hemipteran species, *Nilaparvata lugens*, *Sitobion avenae* and *Acyrthosiphon pisum* (Figure S1B).

Tissue expression profiles showed that *Dc*AKH was expressed at the highest level in the brain, significantly higher than in the ovaries, fat bodies, and midgut in *D. citri* (Figure S2). The levels of *DcAKH* in ovaries of *C*Las-negative and -positive psyllids decreased during the assessment period. However, levels in the ovaries of *C*Las-positive psyllids were significantly higher than those that were *C*Las-negative at all assessment times (Figure 2A). To investigate the role of *DcAKH* in metabolic and reproductive changes induced by *C*Las, we knocked down *DcAKH* expression using RNAi. When fed with ds*DcAKH*, the mRNA level of *DcAKH* in *C*Las-negative and *C*Las-positive ovaries significantly decreased by 89% and 66%, respectively, compared with psyllids fed with ds*GFP* (Figure 2B). Knockdown of *DcAKH* resulted in TAG accumulation and a significant decrease in glycogen (Figures 2C-D). Correspondingly, depletion of *DcAKH* led to a significant increase the size of the lipid droplets (Figure 2E). Compared with the controls, *DcAKH* RNAi slowed ovarian development (Figure 2F), extended the preoviposition period (from 13.3 to 14.2 days in *C*Las-negative psyllids and from 10.7 to 13.4 days in *C*Las-positive psyllids), shortened the oviposition period (from 7.1 to 4.3 days in *C*Las-negative psyllids and from 10.1 to 5.7 days in *C*Las-positive psyllids), and decreased fecundity (from 138.0 to 94.7 days in *C*Las-negative psyllids and from 230.0 to 118.1 days in *C*Las-positive psyllids) (Figures 2G-I). Furthermore, the *C*Las signal and relative titer in the ovaries of *DcAKH* knockdown psyllids significantly decreased (Figures 2J-K). Taken together, it can be concluded that *DcAKH* is involved in *C*Las-induced metabolic and reproductive changes in *D.citri*.

**Figure 2.**
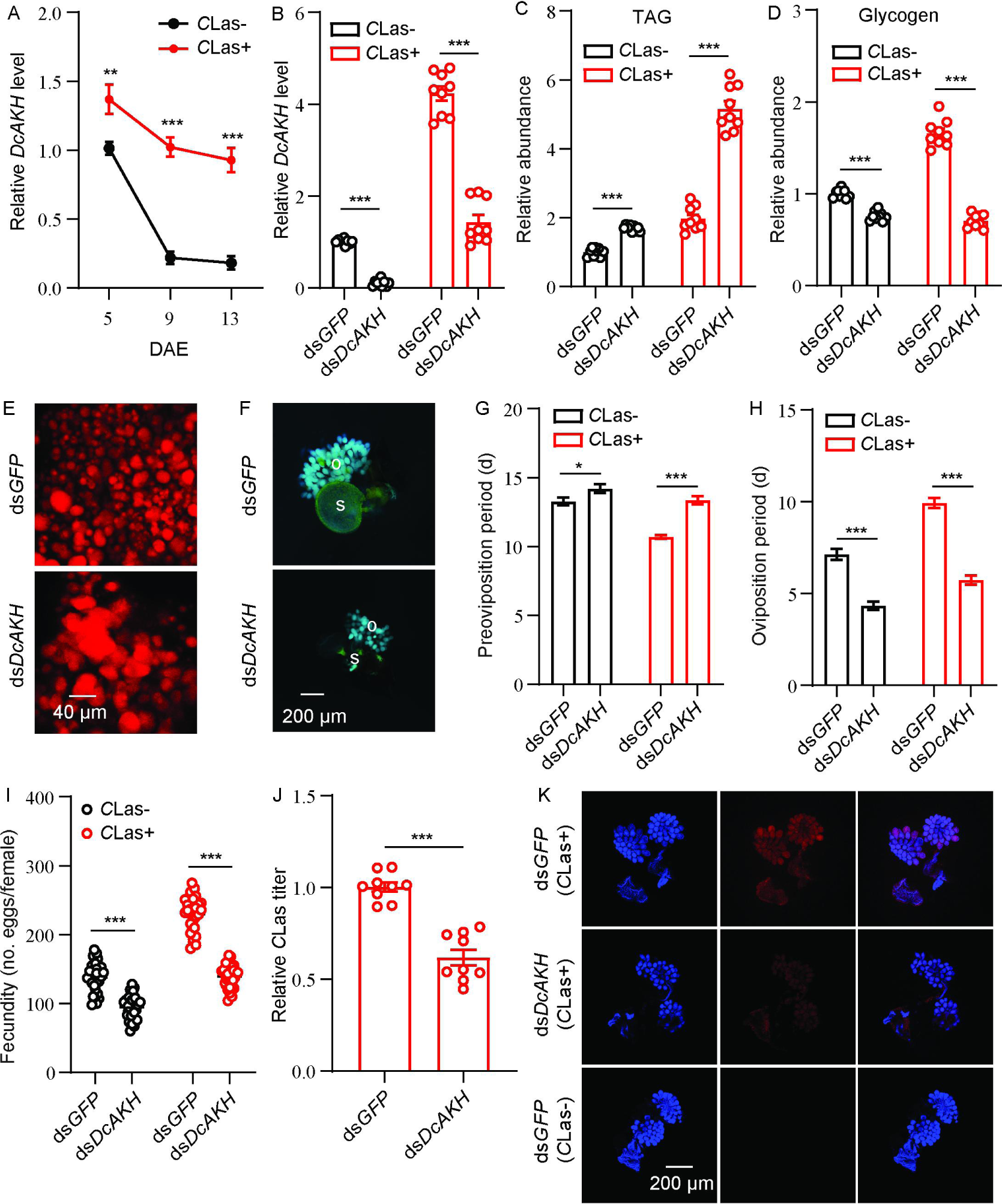
*DcAKH* is involved in the mutualistic relationship between *C*Las and *D. citri* resulting in increased fecundity. (A) Comparison of temporal expression patterns of *DcAKH* between the ovaries of *C*Las- and *C*Las+ psyllids. (B) Efficiency of RNAi of *DcAKH* in *C*Las- and *C*Las+ females treated with ds*DcAKH* for 48 h. (C) Comparison of TAG levels in fat bodies of *C*Las- and *C*Las+ females treated with ds*DcAKH* for 48 h. (D) Comparison of glycogen levels in fat bodies of *C*Las- and *C*Las+ females treated with ds*DcAKH* for 48 h. (E) Lipid droplets stained with Nile red in fat bodies dissected from *C*Las+ females treated with ds*DcAKH* for 48 h. Scale bar = 40 μm. (F) Ovary phenotypes of *C*Las+ females treated with ds*DcAKH* for 48 h. Scale bar = 200 μm. o: ovary, s: spermathecae. (G-I) Comparison of the preoviposition period, oviposition period and the fecundity between *C*Las- and *C*Las+ adults treated with ds*DcAKH*. (J) The *C*Las titer in ovaries of *C*Las+ females treated with ds*DcAKH* for 48 h. (K) Representative confocal images of *C*Las in the reproductive system of *C*Las+ females treated with ds*DcAKH* for 48 h. Scale bar = 200 μm. DAPI: the cell nuclei were stained with DAPI and visualized in blue. *C*Las-Cy3: the *C*Las signal visualized in red by staining with Cy3. Merge: merged imaging of co-localization of cell nuclei and *C*Las. Data are shown as means ± SEM. The significant differences between treatment and controls are indicated by asterisks (Student’s *t*-test, \**p* < 0.05, \*\**p* < 0.01, \*\*\**p* < 0.001).

### *DcAKHR* respond to *DcAKH* and participates in *D. citri*-*C*Las mutualism

The protein tertiary structure of *DcAKHR* was predicted; *DcAKHR* has seven conserved transmembrane (TM1-TM7) domains and is a typical G protein coupled receptor (Figure S3A). A phylogenetic analysis was conducted to evaluate the association of *DcAKHR* with other insect *AKHRs*; *DcAKHR* was most closely related to the *AKHR* of *Nilaparvata lugens* (Figure S3B). To further confirm *Dc*AKH can activate *DcAKHR*, we performed a cell-based calcium mobilization assay. As shown in Figure S3C, *DcAKHR* was strongly activated by *Dc*AKH in a dose-dependent manner, with an EC_50_ value of 2.20 × 10^-5^ mM, but there were no responses when challenged with an empty vector or other evolutionarily-related peptides. In addition, after *DcAKH* knockdown, *DcAKHR* expression in the ovaries significantly decreased (Figure S3D). *DcAKHR* was cloned, sequenced, designated and then used for further studies. Sequence analysis showed that *DcAKHR* consisted of an open reading frame (ORF) of 1290 bp and a 185 bp 3’ UTR. It encodes a putative protein consisting of 429 amino acids with a predicted molecular weight of 48.32 kDa. Tissue expression patterns showed that *DcAKHR* had the highest expression in the midgut, follow by the head, fat bodies, and ovaries (Figure S4). In the ovaries of *C*Las-positive and -negative psyllids, the transcript and protein levels of *DcAKHR* increased over the assessment period with levels of both being higher in *C*Las-positive psyllids than in *C*Las-negative psyllids (Figure 3A). To investigate the role of *DcAKHR* in metabolic and reproductive changes induced by *C*Las, we knocked down *DcAKHR* expression by RNAi. When fed with ds*DcAKHR*, the mRNA levels of *DcAKHR* in the ovaries of *C*Las-negative and *C*Las-positive were significantly decreased by 59.4 and 66.0%, respectively, compared with psyllids fed with ds*GFP*; western blot analysis confirmed that *Dc*AKHR protein levels were also significantly reduced (Figure 3B). Knockdown of *DcAKHR* resulted in TAG accumulation and a significant decrease in glycogen (Figures 3C-D). Correspondingly, depletion of *DcAKHR* led to a significant increase in the size of lipid droplets (Figure 3E). Compared with the controls, *DcAKHR* RNAi interfered with ovarian development (Figure 3F): the preoviposition period was extended (from 13.3 to 16.2 days in *C*Las-negative psyllids and from 10.7 to 12.9 days in *C*Las-positive psyllids); oviposition period was shortened (from 7.1 to 4.6 days in *C*Las-negative psyllids and from 10.1 to 5.3 days in *C*Las-positive psyllids); and fecundity decreased (from 138.0 to 90.6 eggs per female in *C*Las-negative psyllids and from 230.0 to 98.2 eggs per female in *C*Las-positive psyllids). These effects are similar to those caused by *DcAKH* knockdown (Figures 3G-I). Furthermore, the *C*Las signal and relative titer in the ovaries of *DcAKH* knockdown psyllids were significantly decreased (Figures 3J-K). Taken together, *DcAKHR* responds to *DcAKH* and is involved in the *C*Las-induced metabolic and reproductive changes in *D. citri*.

**Figure 3.**
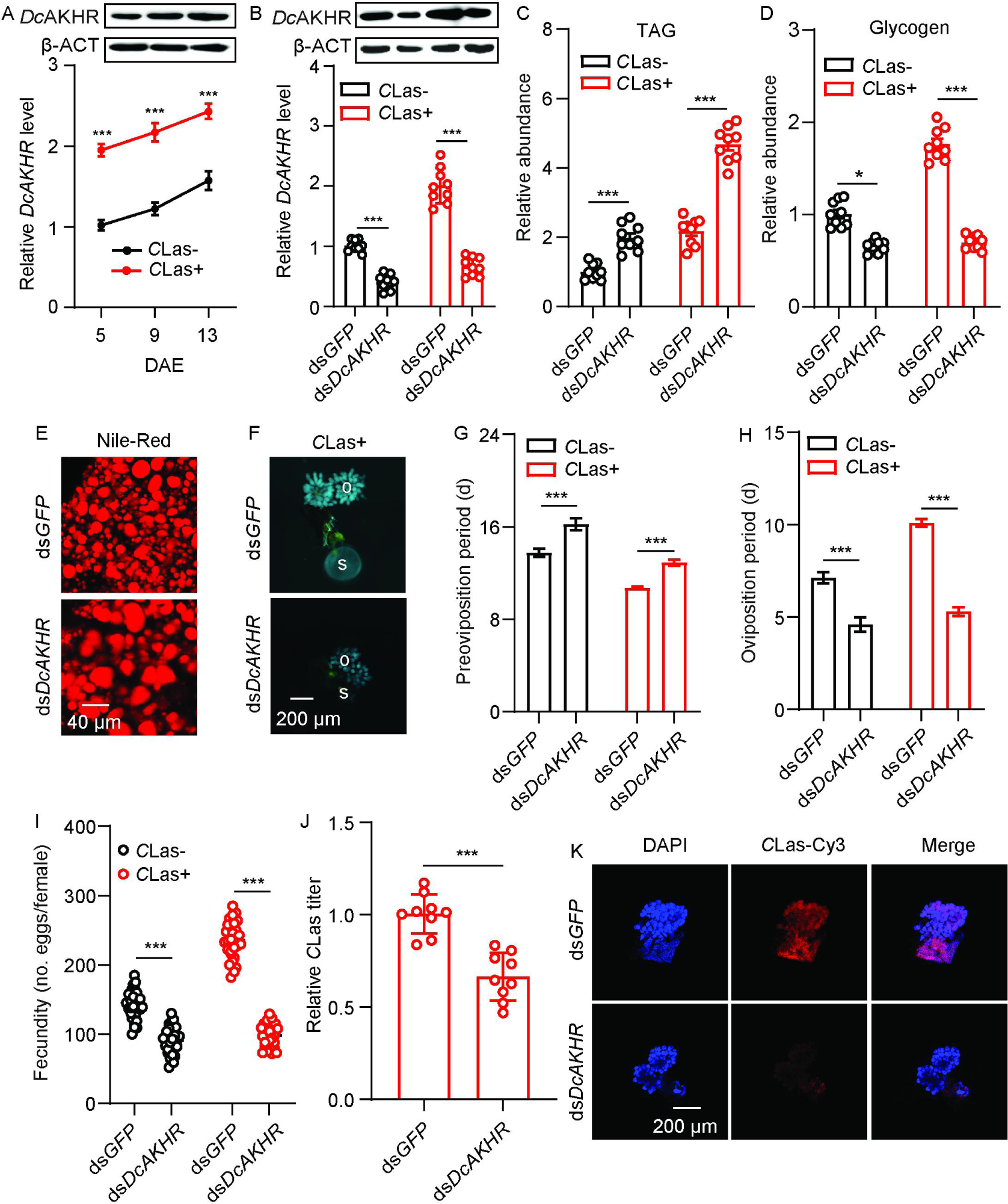
*DcAKHR* is involved in the mutualistic relationship between *C*Las and *D. citri* resulting in increased fecundity. (A) Comparison of temporal expression patterns of *DcAKHR* between the ovaries of *C*Las- and *C*Las+ psyllids. (B) The efficiency of RNAi of *DcAKHR* in *C*Las- and *C*Las+ psyllids treated with ds*DcAKHR* for 48 h. (C) Comparison of TAG levels in fat bodies of *C*Las- and *C*Las+ females treated with ds*DcAKHR* for 48 h. (D) Comparison of glycogen levels in fat bodies of *C*Las- and *C*Las+ females treated with ds*DcAKHR* for 48 h. (E) Lipid droplets stained with Nile red in fat bodies dissected from *C*Las-positive females treated with ds*DcAKHR* for 48 h. Scale bar = 40 μm. (F) Ovary phenotypes of *C*Las+ females treated with ds*DcAKHR* for 48 h. Scale bar = 200 μm. o: ovary, s: spermathecae. (G-I) Comparison of the preoviposition period, oviposition period and the fecundity of *C*Las- and *C*Las+ adults treated with ds*DcAKHR*. (J) The *C*Las titer in the ovaries of *C*Las+ females treated with ds*DcAKHR* for 48 h. (K) Representative confocal images of the reproductive system of *C*Las+ females treated with ds*DcAKHR* for 48 h. Scale bar = 200 μm. DAPI: the cell nuclei were stained with DAPI and visualized in blue. *C*Las-Cy3: the *C*Las signal visualized in red by staining with Cy3. Merge: merged imaging of co-localization of cell nuclei and *C*Las. Data are shown as means ± SEM. The significant differences between treatment and controls are indicated by asterisks (Student’s *t*-test, \*\*\**p* < 0.001).

### miR-34 targeted *DcAKHR*

Based on the small RNA libraries of *D. citri*, we used miRanda and RNAhybrid software to predict the potential miRNAs targeting *DcAKHR*; three putative miRNAs, namely miR-34, miR-2 and miR-14 (Figure 4A), were identified. To confirm the binding activity of these miRNAs with *DcAKHR*, several dual-luciferase reporter assays were conducted. Compared with the controls, the luciferase activity was markedly reduced when miR-34 agomir was co-transfected with the recombinant plasmid containing the full 3’UTR sequence of *DcAKHR*, while miR-2 and miR-14 activities were not significantly changed (Figure 4B). In addition, the reporter activity was recovered when the binding sites of miR-34 were mutated in the *DcAKHR* 3’-UTR (Figure 4C). To determine whether miR-34 specifically targets *DcAKHR*, the following experiments were conducted. Firstly. tissue expression patterns showed that the miR-34 was highly expressed in the head, followed by the fat bodies, ovaries, and midgut (Figure 4D). Secondly, miR-34 has the opposite expression pattern to *DcAKHR* (Fig. 3A), and transcript levels decreased over the assessment period with the levels of both being lower in *C*Las-positive psyllids than in *C*Las-negative psyllids (Figure 4E). Thirdly, after feeding with agomir-34, the relative levels of miR-34 mRNA in the ovaries of *C*Las-negative and *C*Las-positive psyllids increased 1.9- and 2.3-fold, respectively (Figure S5). The mRNA and protein levels of *DcAKHR* were correspondingly increased or decreased after separately feeding psyllids with either antagomir-34 or agomir-34 (Figure 4F). Fourthly, an RNA immunoprecipitation (RIP) assay was performed; *DcAKHR* mRNA was significantly enriched in the anti-AGO-immunoprecipitated RNAs from ovaries of agomir-34-fed female psyllids relative to the control samples (Figure 4G). Taken together, these results strongly suggest that *DcAKHR* is a direct target of miR-34.

**Figure 4.**
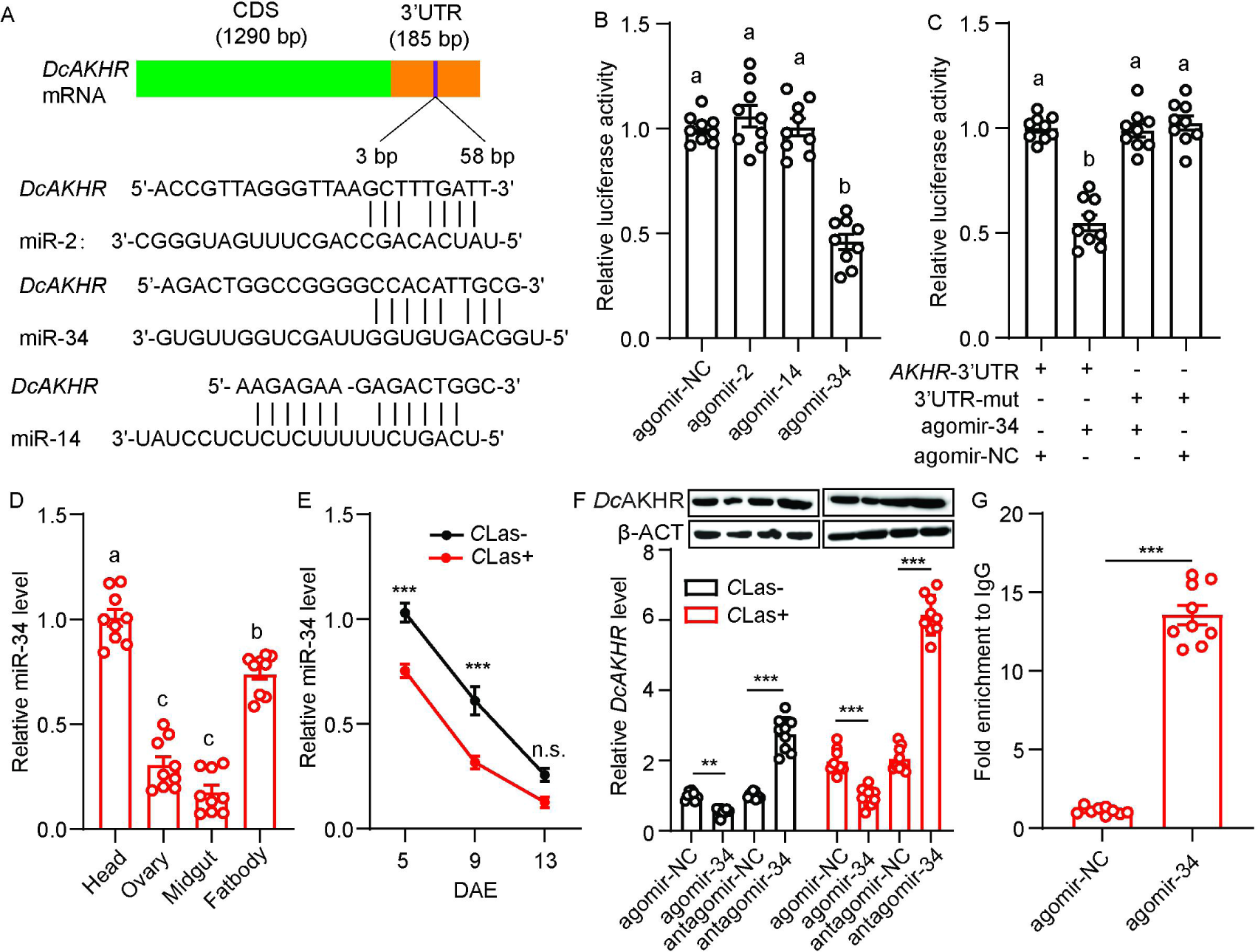
Identification and validation of the target relationship between miR-34 and *DcAKHR.* (A) The putative binding sites of miRNAs in the *DcAKHR* 3’-UTR as predicted by miRanda and RNAhybrid. (B) Dual-luciferase reporter assays using HEK293T cells co-transfected with miRNA agomir and recombinant pmirGLO vectors containing the predicted binding sites for miR-2, miR-14 and miR-34 in the CDS of *DcAKHR.* (C) Dual-luciferase reporter assays using HEK293T cells co-transfected with miR-34 agomir plus recombinant pmirGLO vectors containing *DcAKHR-*3’UTR or mutated *DcAKHR-*3’UTR. (D) Tissue expression pattern of miR-34 in *C*Las+ female adults at 7 DAE in the head, ovary, fat body, and midgut. (E) Comparison of temporal expression patterns of miR-34 in ovaries of *C*Las- and *C*Las+ females. (F) Effect of miR-34 agomir and antagomir treatments on *Dc*AKHR mRNA expression and protein level in ovaries of *C*Las- and *C*Las+ psyllids after 48 h. (G) Relative expression of miR-34 targeted *DcAKHR in vivo* as demonstrated by an RNA immunoprecipitation assay. Data are shown as mean ± SEM. For B-D, significant differences among the different treatments are indicated by lowercase letters above the bars (one-way ANOVA followed by Tukey’s Honestly Significant Difference test at *P* < 0.05). The significant differences between treatment and control are indicated by asterisks in E-G (Student’s *t*-test, \*\**p* < 0.01, \*\*\**p* < 0.001).

### miR34 participates in *D. citri*-*C*Las mutualism in ovaries

To investigate the roles of miR-34 in the interaction between *D. citri* and *C*Las, *C*Las-positive and –negative females were fed with agomir-negative control (NC) or with miR-34 agomir. After feeding with miR-34 agomir, there was a significant accumulation of TAG and a decrease in glycogen levels in the two populations (Figures 5A-B). The size of lipid droplets in the fat bodies of *C*Las-positive psyllids were bigger than those of the control (Figure 5C). After treatment with miR-34 agomir, ovarian development of *C*Las-positive psyllids was reduced (Figure 5D). In addition, when *C*Las-negative and *C*Las-positive psyllids were fed with miR-34 agomir, the oviposition period of both psyllid types was shortened, preoviposition period was markedly extended, and fecundity significantly decreased compared with the relevant control groups (Figures 5E-G); these phenotypes mimic those expressed after treatment with ds*DcAKHR*. Moreover, the *C*Las signals and relative titer in *C*Las-positive ovaries were significantly reduced (Figures 5H-I). All the results indicate that miR-34 suppresses *DcAKHR* expression and is involved in the mutualistic interaction.

**Figure 5.**
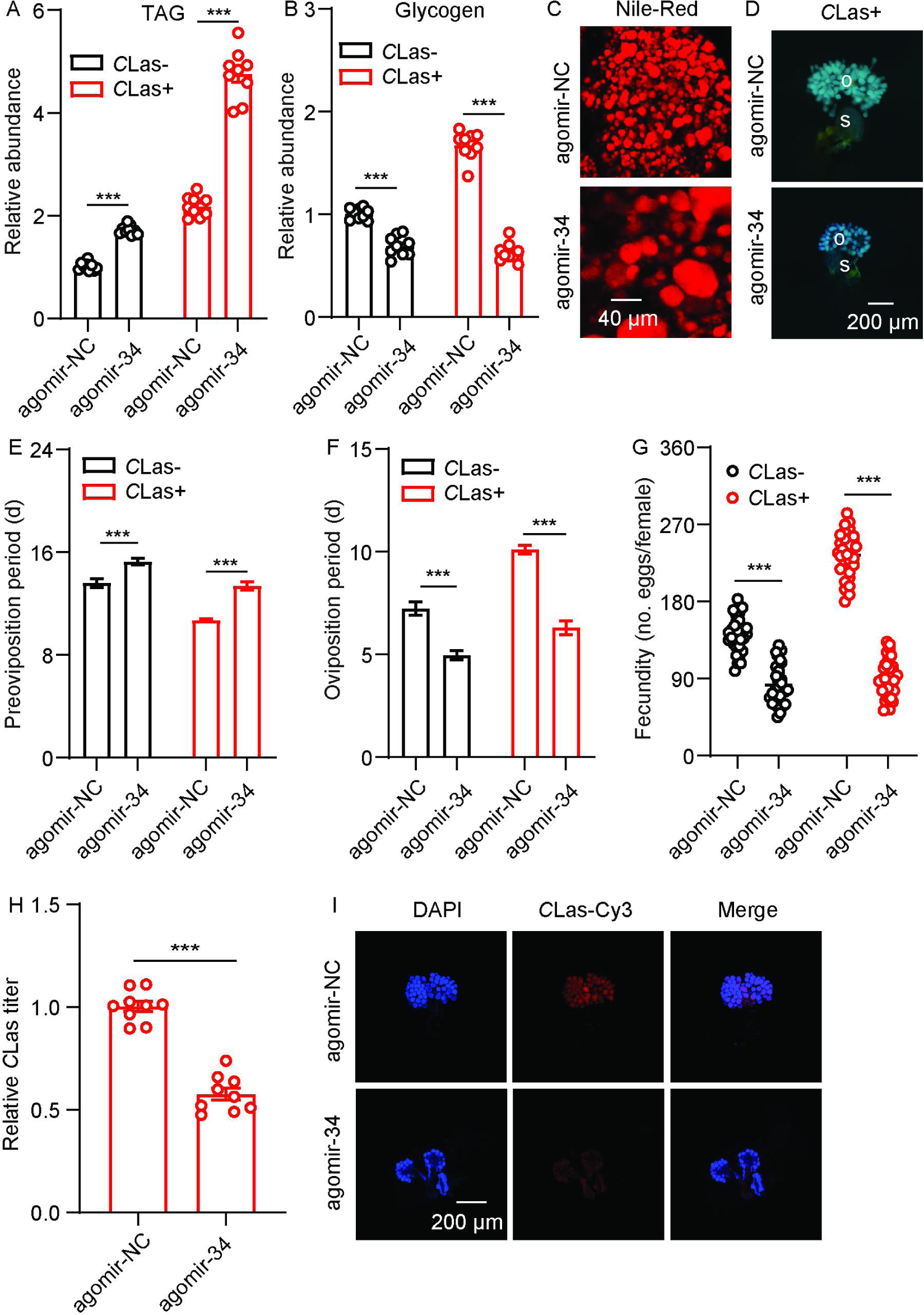
miR-34 participation in mutualistic interactions between *D. citri* and *C*Las. (A) Comparison of TAG levels in fat bodies of *C*Las- and *C*Las+ females treated with agomir-34 for 48 h. (B) Comparison of glycogen levels in the fat bodies of *C*Las- and *C*Las+ females treated with agomir-34 for 48 h. (C) Lipid droplets stained with Nile red in fat bodies dissected from *C*Las+ females treated with agomir-34 for 48 h. Scale bar = 40 μm. (D) Ovary phenotypes of *C*Las+ female treated with agomir-34 for 48 h. Scale bar = 200 μm. o: ovary, s: spermathecae. (E-G) Comparison of the preoviposition period, oviposition period and the fecundity between *C*Las- and *C*Las+ adults treated with agomir-34. (H) *C*Las titer in ovaries of *C*Las+ females treated with agomir-34 for 48 h. (I) Representative confocal images of *C*Las in the reproductive system of *C*Las+ females treated with agomir-34 for 48 h. Scale bar = 200 μm. The signals of DAPI and *C*Las-Cy3 are same as described in Figure 2. Data are shown as means ± SEM. The significant differences between treatment and controls are indicated by asterisks (Student’s *t*-test, \*\*\**p* < 0.001).

### The JH signaling pathway is regulated by the AKH pathway and is involved in the increase in fecundity induced by *C*Las

To evaluate whether the JH signaling pathway is regulated by the AKH signaling pathway, we assayed the relative JH titers and the expression levels of several JH pathway-related genes in *DcAKH-*deficient, *C*Las-positive females. Compared to control females, the relative JH titer in the abdomen decreased significantly after *dsDcAKH* treatment for 48 h (Figure 6A). After feeding with *dsDcAKH* for 48 h, the expression levels of the JH receptor gene, *DcMet,* and the downstream transcription factor, *DcKr-h1*, of the JH signaling pathway were significantly reduced in the fat bodies and ovaries (Figures 6B-C). In addition, the expression levels of *DcVg1-like*, *DcVgA1-like* and *DcV*gR, three important downstream genes of the JH signaling pathway related to ovarian development were lower in *DcAKH-*deficient *C*Las-positive females than in controls after dsRNA feeding for 48 h (Figures 6B-C). Similar results were observed in the *DcAKHR-*deficient *C*Las-positive females (Figures 6D-F) as well as *C*Las-positive females fed with agomir-34 (Figures 6G-I). These results suggest that JH signaling pathway is regulated by the AKH-AKHR-miR-34 signaling pathway and is involved in the increased fecundity of *D. citri* induced by *C*Las.

**Figure 6.**
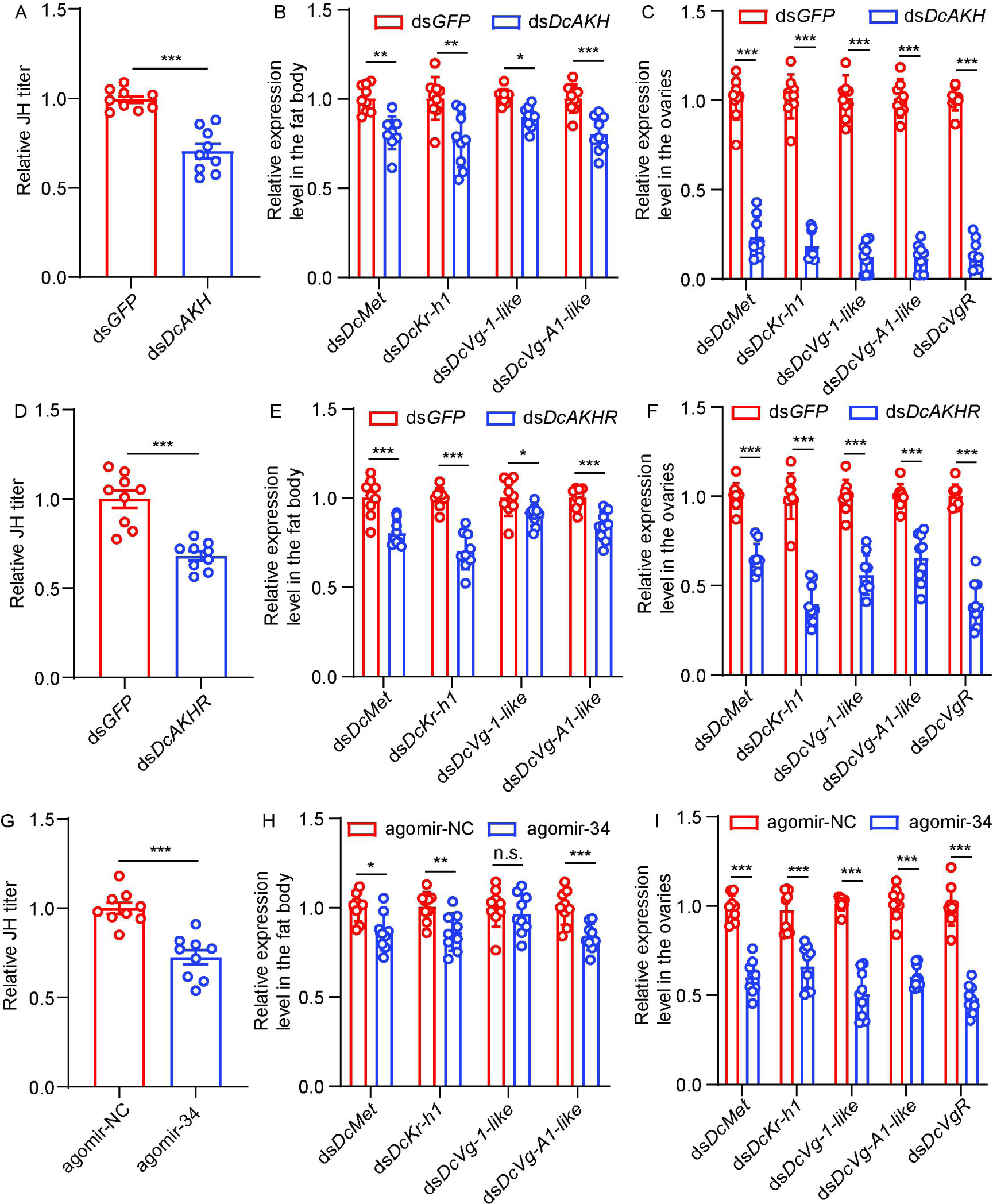
The JH signaling pathway is regulated by AKH signaling pathway and is involved in the increase in fecundity of *D. citri* induced by *C*Las. (A) JH titer in the abdomen of *C*Las+ females treated with ds*DcAKH* for 48 h. (B) Effects of ds*DcAKH* treatment on mRNA level of JH signaling pathway in fat bodies of *C*Las+ females. (C) Effects of ds*DcAKH* treatment on mRNA levels of components of the JH signaling pathway in the ovaries of *C*Las+ females. (D) JH titers in the abdomens of *C*las+ females treated with ds*DcAKHR* for 48 h. (E) Effects of ds*DcAKHR* treatment on mRNA level of JH signaling pathway in fat bodies of *C*Las+ females. (F) Effects of ds*DcAKHR* treatment on mRNA levels of components of the JH signaling pathway in ovaries of *C*Las+ females. (G) JH titer in abdomen of *C*Las+ females treated with agomir-34 for 48 h. (H) Effects of agomir-34 treatment on mRNA levels of components of the JH signaling pathway in fat bodies of *C*Las+ females. (I) Effects of agomir-34 treatment on mRNA levels of components of the JH signaling pathway in the ovaries of *C*Las+ females.

## Discussion

An increasing number of studies have focused on the effects of vector-virus interactions on reproduction. For example, rice gall dwarf virus is transmitted vertically by hitchhiking on insect sperm, thereby promoting long-term viral epidemics and persistence in nature [2]. Berasategui et al. (2022) [3] found a mutualistic relationship between *Chelymorpha alternans* and *Fusarium oxysporum*, where the pupae were protected in exchange for the dissemination and propagation of the fungus. Although there is limited research on the mechanisms underlying vector-bacteria interactions. In *D. citri*-*C*Las interaction, *C*Las operates host hormone signaling and miRNA to mediate the mutualistic interactio between *D. citri* fecundity and its replication [35]. Singh and Linksvayer (2020) [41] found that *Wolbachia*-infected colonies of *Monomorium pharaonis* exhibited increased colony-level growth, accelerated colony reproduction, and shortened colony life cycles compared to uninfected colonies.

The fat body is the major insect storage organ that produces the majority of hemolymph-born vitellogenic proteins and critically contributes to insect ovary development [16]. In this study, after infection with *C*Las, the preoviposition period of *D. citri* was shortened, oviposition period was prolonged, and fecundity was significantly increased. These results suggests that there is a mutualistic interaction in *D. citri* ovaries with *C*Las. *C*Las accelerates ovarian development and prolongs the oviposition period thereby enhancing the fecundity of *D. citri*; in turn, *D. citri* provides sites and nutrients for *C*Las replication during ovarian development. *C*Las-positive *D. citri* require more energy to support this increased fecundity. Hence, the TAG and glycogen levels of both *C*Las-positive and *C*Las-negative were assayed. After infection with *C*Las, the TAG and glycogen levels significantly increased, and the average size of lipid droplets in the fat bodies were distinctly larger, indicating that the increase in lipid mobilization may prepare for an increase in fecundity.

The total amount of lipids present in fat bodies is a balance between lipogenesis and lipolysis [42], and both processes are essential for lipid homeostasis. Disruption of either lipogenesis or lipolysis could interfere with nutritional and physiological conditions in fat bodies and affect vitellogenesis. AKH/AKHR mediated lipolytic systems are essential for lipid provision and nutrient transfer during reproduction. In *Gryllus bimaculatus* [43,44] and *Caenorhabditis elegans* [45], injections of AKH interfered with egg production by inhibiting the production of vitellogenins and the accumulation of lipid stores. In *Bactrocera dorsalis*, AKHR knockdown led to decreased lipolytic activity and delayed oocyte maturation [29] and these authors suggested the decline in fecundity may be due to the inability to utilize the lipid stores of fat bodies to promote oocyte maturation. In *Glossina morsitans*, the silencing of *AKHR* and *brummer* led to the inability to mobilize lipid reserves in fat bodies during pregnancy, slowed down the development of oocytes, and caused embryogenesis to fail, indicating that female fertility depends on lipid metabolism regulated by AKHR [24]. These results suggest that lipid homeostasis appears critical for insect fecundity, as its disruption substantially suppresses egg production and negatively affects vitellogenesis.

In the study, *DcAKH* and *DcAKHR* played important roles in the increased lipid metabolism and fecundity of *D. citri* mediated by *C*Las. The cDNA sequences encoding AKH and AKHR from *D. citri* were cloned and identified, and phylogenetic analysis showed that *DcAKH* and *DcAKHR* were orthologous to other hemipteran AKHs and AKHRs. Functional analysis showed that *DcAKHR* shares properties found with other AKHRs. *DcAKHR-*transfected CHO cells could be activated by AKH peptide, while it did not response to Crz peptide, despite the sequences and gene structures of AKH and Crz being closely related [21]. *DcAKH* and *DcAKHR* were more highly expressed in the ovaries of *C*Las-positive psyllids than those of *C*Las-negative individuals. Knockdown of *DcAKH* and *DcAKHR* resulted in TAG accumulation and a significant decrease in glycogen levels in the fat bodies. Most importantly, the *C*Las titer in the ovaries of *C*Las-positive psyllids and fecundity significantly reduced when feeding with *dsDcAKH* and *dsDcAKHR*. Our results support the idea that reproductive disruption from AKH/AKHR knockdown is due to the inability of female to mobilize lipid reserves in the fat bodies required for oocyte maturation during vitellogenesis.

To date, there are no reports on the post-transcriptional regulation of AKHR. As important post-transcriptional regulators, miRNAs generally suppress their target genes by triggering mRNA degradation or translational repression [32]. There is increasing evidence implicating miRNAs in the metabolic processes of insects, particularly in relation to reproduction. In addition to functions of miR-277 described in the Introduction, this miRNA targets ilp7 and ilp8 thereby controlling lipid metabolism and reproduction in *A. aegypti* [40]. miR-8 depletion in mosquitoes resulted in an increase of its target gene, *secreted wingless-interacting molecule*, and in lipid accumulation in developing oocytes [46]. Depletion of miR-8 in mosquitoes by antagomir injection upregulates *secreted wingless-interacting molecule* and decreases the level of lipid in ovaries to block vitellogenesis post blood meals [47]. A switch from catabolic to anabolic amino acid metabolism in fat bodies mediated by miR-276 restricts mosquito investment into oogenesis and benefits the development of *Plasmodium falciparum* [48]. In the current study, based on *in vitro* and *in vivo* experiments, we found that host miR-34 targets *DcAKHR*. miR-34 was observed to have an opposite expression trend to *DcAKHR*, and its expression in the ovaries of *C*Las-positive psyllids was lower than in those of CLas-negative individuals. Treatment of miR-34 agomir and antagomir, respectively, resulted in a significant decrease and increase in *DcAKHR* mRNA and protein levels. In addition, miR-34 overexpression not only led to the accumulation of TAG and the reduction of glycogen levels in fat bodies, but also decreased fecundity and *C*Las titers in the ovaries similar to the phenotypes caused by silencing of *DcAKHR*. This indicates that *C*Las inhibited host miR-34 to enhance *DcAKHR* expression and improve the lipid metabolism and fecundity. This study is the first to report the manipulation of host miRNAs and target genes by a bacterium to affect lipid metabolism and reproduction of their vector.

Vitellogenesis is strongly linked with insect JH production [49]. The fat body is an important tissue that senses and integrates various nutritional and hormonal signals required for the regulation of vitellogenesis [16]. Moreover, JH is also involved in the nutrient-dependent regulation of insect reproduction, but the mechanism of action remains obscure. Lu et al. (2016a, b) [50,51] have demonstrated that nutritional signaling during female reproduction induces JH biosynthesis that in turn stimulates vitellogenin expression in fat bodies and egg production in *N. lugens.* In order to evaluate the possible relationship between AKH/AKHR-mediated lipolysis and JH-dependent vitellogenesis, the expressions of JH signaling pathway-related genes in fat bodies and ovaries were determined after knockdown of *DcAKH* and *DcAKHR*. We found that reducing *DcAKH* or *DcAKHR* by RNAi in *C*Las-positive *D. citri* not only decreased JH titer but also resulted in a decrease in the expression of *DcMet* and *DcKr-h1* in the fat bodies and ovaries, as well as the egg development related genes, *DcVg-1-like*, *DcVg-A1-like* and *DcVgR*. Similar results were detected in the agomir-34 treatment. Our results suggest that AKH/AKHR-based lipolysis plays an important regulatory role in the JH signaling pathway during *D. citri-C*Las interactions.

In conclusion, we proposed the scheme of events following *C*Las infection presented in Figure 7. Upon infection with *C*Las, *D. citri* exhibits enhanced fecundity compared to uninfected individuals.The interaction between *C*Las and *D. citri* affecting reproduction is a win-win strategy; the increased offspring of *D. citri* contributes to a higher presence of *C*Las in the field. *C*Las upregulates the AKH/AKHR signaling and downregulates miR-34 to increase lipid metabolism and activate JH-dependent vitellogenesis, thereby improving the fecundity of *C*Las-positive females. Overall, our results not only broaden our understanding of the molecular mechanism underlying the *D. citri-C*Las interaction, but also provide valuable insights for further integrated management of *D. citri* and HLB by targeting the *DcAKH/DcAKHR* signaling pathway.

**Figure 7.**
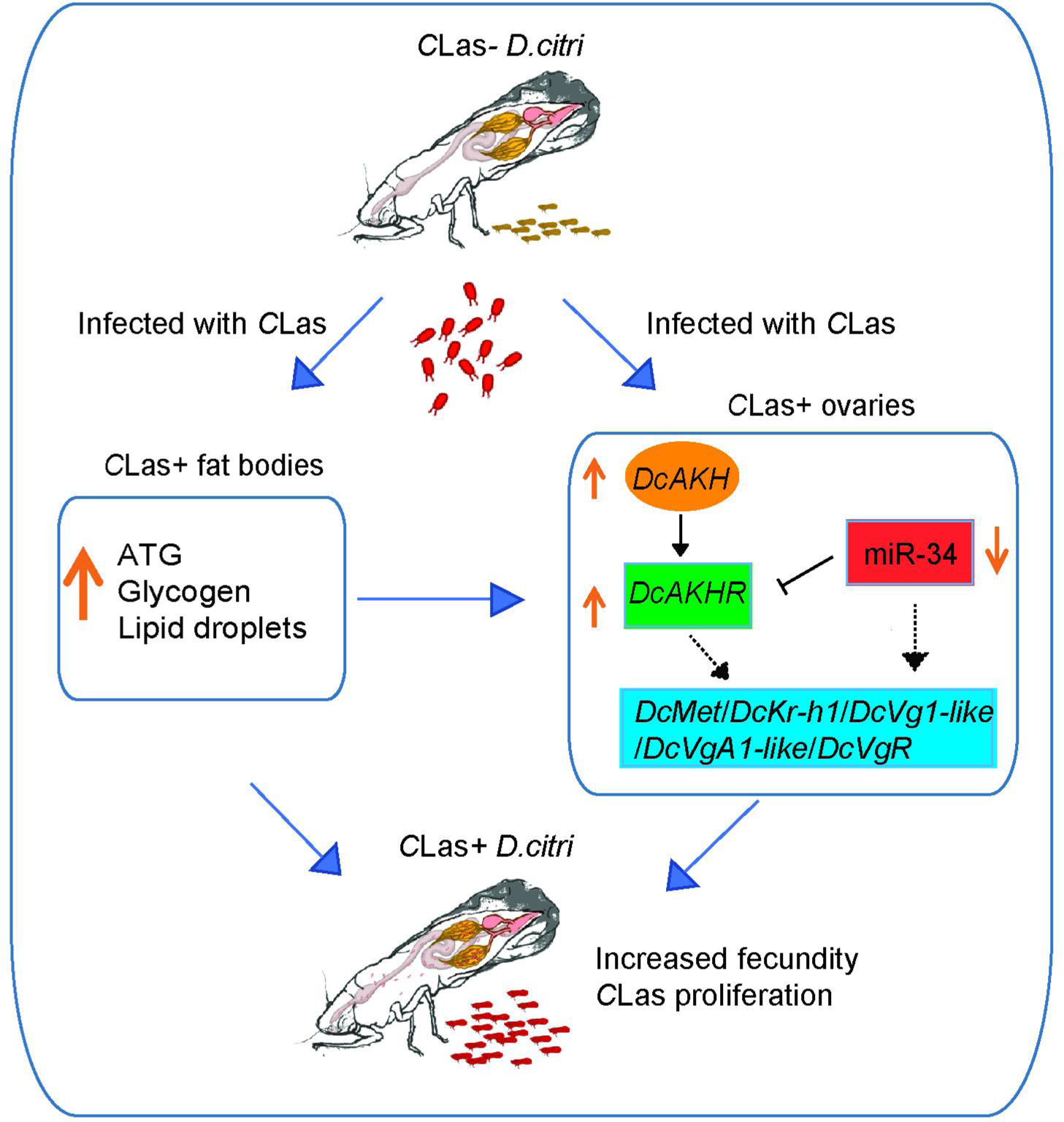
Mechanisms linking metabolism and reproduction of *D. citri* induced by *C*Las. After infection with *C*Las, the TAG and glycogen levels in fat bodies of *C*Las-positive psyllids significantly increased as well as the size of lipid droplets. In ovaries, *C*Las upregulates the AKH/AKHR signaling and downregulates miR-34 to increase lipid metabolism and activate JH-dependent vitellogenesis, thereby improving the fecundity of *C*Las-positive females. *D. citri* are more fecund than their uninfected counterparts. The interaction between *C*Las and *D. citri* affecting reproduction is a win-win strategy; the more offspring of *D. citri*, the more *C*Las in the field.

## Materials and Methods

### Host plants and insect colonies

The *C*Las-negative (healthy) and *C*Las-positive *D. citri* colonies were obtained from a laboratory culture continuously reared on the healthy and *C*Las-infected lemon (*Citrus* ×*limon* (L.) Osbeck) plants. Monthly monitoring of the *C*Las infection in both the lemon plants and psyllids was conducted using quantitative polymerase chain reaction (qPCR). The two populations were raised in different incubators with same conditions (26 ± 1 ℃, 65 ± 5% relative humidity (RH) and a 14-h light: 10-h dark cycle). *C*Las-negative and *C*Las-positive lemon plants were grown in two separate glasshouses.

### Bioinformatics and phylogenetic analyses

The physicochemical properties of *DcAKH* (GenBank accession number: MG550150.1) and *DcAKHR* (GenBank accession number: OR259432) were analyzed using the online bioinformatics ProtParam tool (http://web.expasy.org/protparam/). Search and download amino acid sequences of other species in the NCBI database. Homologous protein sequences of AKH and AKHR were aligned with those of other insects using DNAman6.0.3 software. Phylogenetic trees were constructed with the neighbor-joining method in MEGA5.10 software, with 1000 bootstrap replicates; bootstrap values > 50% shown on the tree.

### Assays of reproductive parameters

Tested females from two colonies were selected and paired with healthy males. One female and one male were placed onto young vegetative shoots (flush) of healthy lemon plants to promote oviposition; the flush and insects were sealed with tied white mesh bag (15 × 20 cm). These lemon plants were held in an incubator (25 ± 1 ℃, 65 ± 5% RH, 14L: 10D photoperiod). After 24h, the preoviposition period and oviposition period were recorded, the number of eggs laid per female were counted, and this pair of psyllids were transferred to new flush to continue egg laying. Eggs were counted daily until all females died. All the experiments were performed using three replications, and about 15 pairs of *D. citri* for each replication.

### Quantitative RT-PCR (qRT-PCR)

For mRNA expression analysis, total RNA was extracted using TRIzol reagent (Invitrogen, Carlsbad, CA, United States), and the synthesis of first-strand cDNA was performed using a PrimeScript™ II 1st Strand cDNA Synthesis Kit (Takara, Beijing, China) in accordance with the manufacturer’s instructions. qRT-PCR was carried out by using the TB Green® Premix Ex Taq™ II (Takara, Beijing, China) on an ABI PRISM® 7500 Real-Time System (Applied Biosystems, Foster City, CA, USA). The beta-actin (*Dcβ-ACT*, GenBank XM_026823249.1) gene was used as an internal control to normalize gene expression levels. For miRNA expression analysis, miRNA was extracted using a miRcute miRNA Isolation Kit (TIANGEN, Beijing, China), synthesized using a miRcute Plus miRNA First-Strand cDNA Kit (TIANGEN, Beijing, China), and quantified using the miRcute Plus miRNA qPCR Kit (SYBR Green) (TIANGEN, Beijing, China). U6 snRNA was used as an internal control to normalize the expression of miRNA. All the primers used are listed in Supplementary Table 1.

### Luciferase activity assay

The 185 bp sequence of the 3’UTR containing the predicted target sites for miR-34 in *DcAKHR* was cloned into the pmirGLO vector (Promega, Wisconsin, USA) downstream of the luciferase gene to construct the recombinant plasmid, *DcAKHR*-3’UTR-pmirGLO, using the pEASY®-Basic Seamless Cloning and Assembly Kit. The mutated 3’UTR sequence was amplified and cloned into the pmirGLO vector to generate the *DcAKHR*-3’UTR mutant-pmirGLO plasmid. The agomir and antagomir of miRNAs were chemically synthesized and modified by GenePharma (Shanghai, China) with chemically modified RNA oligos of the same sequence or anti-sense oligonucleotides of the miRNA. The negative control for agomir and antagomir was provided by the manufacturer. According to the manufacturer’s instructions, the constructed vector (500 ng) and miRNA agomir (275 nM) were co-transferred into HEK293T cells in a 24-well plate using a Calcium Phosphate Cell Transfection Kit (Beyotime, Nanjing, China). After co-transfection for 24 h, the activities of firefly and Renilla luciferase were detected using the Dual-Glo Luciferase Assay System (Promega, Madison, WI, USA), and the average luciferase activity was calculated.

### Fluorescence in situ hybridization (FISH)

FISH using a *C*Las probe was carried out as previously described [52]. Under a dissecting microscope, the ovaries of *C*Las-negative and *C*Las-postive *D. citri* were dissected in 1 × phosphate-buffered saline (PBS) (Beyotime Biotechnology, Shanghai, China). Firstly, the isolated ovaries were fixed in Carnoy’s fixative (glacial acetic acid-ethanol-chloroform, 1: 3: 6, vol/vol) for 12 h at 25 ℃, washed four times (5 min each time) with 6% H_2_O_2_ in 80% ethanol and then three times (10-15 min per time) with PBST (1 × PBS: TritonX-100, 99.7: 0.3, (vol/vol)). Secondly, the samples were pre-incubated three times (10 min per time) in hybridization buffer (20 mM Tris-HCl, pH 8.0, 0.9 M NaCl, 30% formamide (vol/vol), 0.01% sodium dodecyl sulfate (wt/wol)) containing 10 pmol/mL of each probe for 24 h at 25 ℃, then washed three times with PBST. Thirdly, nuclei were stained with 0.1 mg/mL of 40, 60-diamidino-2-phenylindole (DAPI) (Sigma-Aldrich, St Louis, MO, USA) for 15 min, then washed once with PBST. Finally, the stained ovaries were mounted in mounting medium. The slides were scanned under a Leica TCS-SP8 (Leica Microsystems Exton, PA USA) confocal microscope; excitation lasers emitting at 405 nm and 550 nm were used to detect the DAPI and Cy3 signals, respectively, and sequential scanning was used to avoid the signal overlap of probes. Image processing was completed using Leica LAS-AF software (v2.6.0). Specificity of detection was carried out using without the probe and *C*Las-negative controls. Three FISH tests were performed and at least 20 ovaries were viewed under the microscope to confirm the repeatability. The probe sequences used in the current study are listed in Supplementary Table 1.

### Western blotting

The total proteins from *D. citri* ovaries were extracted using RIPA protein lysis buffer (50 mM Tris pH 7.4, 1% Triton X-100, 150 mM NaCl, 1% sodium deoxycholate and 0.1% SDS) with 1 mM PMSF. Proteins concentrations were quantified using a BCA protein assay kit (Beyotime, Jiangsu, China). Aliquots of 60 μg protein were separated by 12% SDS-PAGE and transferred to polyvinylidene fluoride membranes (Millipore). Subsequently, the membranes were incubated with the primary antibody against *DcAKHR* (1: 1000 dilution, ABclonal Technology Co., Ltd., Wuhan, China) for 12 h at 4 °C and then with the secondary antibody (goat anti-rabbit IgG conjugated with HRP, 1: 10000 dilution) for 2 h at 25 °C. A mouse monoclonal antibody against β-actin (TransGen Biotech, Beijing, China) was used as a control. Immunoreactivity was imaged with the enhanced chemiluminescence with Azure C600 multifunctional molecular imaging system.

### Nile red staining

The fat bodies were dissected from adult females 5 DAE, fixed in 4% paraformaldehyde for 30 min at 25 ℃, and washed twice with 1 × PBS. Lipid droplets were incubated for 30 min within the mixture of Nile red (0.1 μg/μL, Beijing Coolaber Technology Co., Ltd) and DAPI (0.05 μg/μL), and then washed twice with 1 × PBS. The samples were imaged using a laser scanning microscopy (TCS-SP8, Leica Microsystems Exton, PA USA).

### Determination of TAG levels

TAG levels were measured using a Triglycerides Colorimetric Assay (Cayman) in accordance with the manufacturer’s instructions. In short, thirty fat bodies were homogenized in 100 μL of Diluent Assay Reagent. Then 10 μL of supernatant was incubated with Enzyme Mixture solution, the TAG contents from the measurements were normalized to protein levels in the supernatant of the samples determined using a BCA protein assay (Thermo Fisher Scientific). Three independent biological replicates were analyzed for each treatment, and each treatment included three technical replicates.

### Determination of glycogen levels

Glycogen levels were assayed using a Glycogen Assay Kit (Cayman) in accordance with the manufacturer’s instructions. In brief, thirty fat bodies were homogenized in 100 μL of Diluent Assay Reagent, and 10 μL of supernatant was incubated with Enzyme Mixture solution and Developer Mixture. The glycogen levels from the measurements were normalized to protein levels in the supernatant of the sample determined using a BCA protein assay (Thermo Fisher Scientific). Three independent biological replicates were analyzed for each treatment, and each treatment included three technical replicates.

### dsRNA synthesis and RNAi assay

dsRNAs of *DcAKH* and *DcAKHR* were transcribed by a Transcript Aid T7 High Yield kit (Thermo Scientific, Wilmington, DE, United states) and purified using the GeneJET RNA Purification kit (Thermo Scientific). miRNA agomir, agomir negative control, miRNA antagomir, and antagomir negative control were synthesized in the Shanghai GenePharma Co. Ltd. (Shanghai, China). The RNAi and miRNA treatments were performed by feeding dsRNA or miRNA antagomir/agomir through an artificial diet as described previously [53]. Briefly, twenty females at 7 DAE were placed into a glass cylinder (25 × 75 mm) and sealed with two stretched paramembranes. 200 μL of 20% (w:v) sucrose mixed with dsRNA was placed between two paramembranes for feeding. The final ingestion concentrations of ds*DcAKH* and ds*DcAKHR* were 200 ng/μL and 300 ng/μL, respectively. After feeding with dsRNA for 48 h, the treated females were assayed as follows. The ovaries were dissected or qRT-PCR and western blot analysis. Reproductive parameters including the pre-oviposition period, oviposition period, and fecundity were recorded. *C*Las titers in the ovaries were assayed. Ovary morphologies were observed using a Ultra-Depth Three-Dimensional Microscope (VHX-500). Fat bodies were stained with Nile red. TAG levels and glycogen levels were determined. Green fluorescent protein (GFP) was used as a reporter gene. Feeding ds*GFP* was used as the control when psyllids were fed with dsRNA. All the experiments were performed in three replications, and each replication included 15-20 pairs of *D. citri*.

### Heterologous expression and calcium mobilization assay

The complete ORF of *DcAKHR* was cloned into the expression vector pcDNA3.1^+^; cloning was confirmed by sequencing (TSINGKE Bio). Endotoxin-free plasmid DNA of the vector was extracted, and all tested peptides were synthesized by Sangon Biotech (Shanghai, China) at a purity >95 % (see Fig. 1B for peptide structures). Chinese hamster ovary (CHO-WTA11) cells supplemented with Gα16 subunit and aequorin were used [54] and transfected with pcDNA3.1^+^-*DcAKHR* or empty pcDNA3.1^+^ (negative control) vectors using Lipofectamine 2000 (Thermo Fisher Scientifc, Waltham, MA USA). The calcium mobilization assay was performed with slight modification as described by Gui et al. (2017) [55] and Shi et al. (2017) [56]. In brief, the cells were incubated for 3 h in the dark with coelenterazine h (Promega, Madison, WI, USA) 48 h after the transfection. Then, serial dilutions of peptide ligands (10^−6^ to 10^1^ μM) were loaded into opaque 96-well plates. Luminescence was measured over 15 s using a SpectraMax i3x Multi Mode Microplate Reader (Molecular Devices). Medium alone was used as a blank control and 100 μM ATP was served as a positive control. Each experiment was repeated three times.

### JH titer detection

JH titers were assessed using the Insect JH Ⅲ ELISA kit (Shanghai Enzyme-linked Biotechnology Co., Ltd. Shanghai, China) following the manufacturer’s protocol.

### Statistical analysis

All statistical values are presented as means ± SEM. Statistical analyses were performed by GraphPad Prism 8.0 software. Student’s *t*-test was used to determine the pairwise comparisons at the following significance levels (\**p* < 0.05, \*\**p* < 0.01, \*\*\**p* < 0.001). For multiple comparisons, one-way ANOVA with Tukey’s Honest Significant Difference tests used to separate means at *p* < 0.05.

## Acknowledgments

This research was supported by the National Natural Science Foundation of China, grant number 32102193, and the Open Competition Program of Ten Major Directions of Agricultural Science and Technology Innovation for the 14th Five-Year Plan of Guangdong Province, grant number 2022SDZG07.

## Author Contributions

Conceptualization: Jiayun Li, Paul Holford, George Andrew Charles Beattie, Yurong He, Yijing Cen, Xiaoge Nian.

Data curation: Jiayun Li, Shujie Wu, Jielan He, Xiaoge Nian.

Formal analysis: Jiayun Li, Xiaoge Nian, Desen Wang.

Funding acquisition: Xiaoge Nian, Yurong He, Yijing Cen.

Investigation: Jiayun Li, Shujie Wu, Jielan He, Shijian Tan, Xiaoge Nian.

Methodology: Jiayun Li, Paul Holford, George Andrew Charles Beattie, Yurong He, Xiaoge Nian, Yijing Cen.

Project administration: Xiaoge Nian, Yijing Cen.

Resources: Jiayun Li, Shujie Wu, Jielan He, Xiaoge Nian, Yijing Cen.

Software: Jiayun Li, Xiaoge Nian, Desen Wang.

Supervision: Yurong He, Yijing Cen.

Validation: Jiayun Li, Paul Holford, George Andrew Charles Beattie, Xiaoge Nian.

Visualization: Jiayun Li, Paul Holford, George Andrew Charles Beattie, Yurong He, Xiaoge Nian, Yijing Cen.

Writing-original draft: Jiayun Li, Xiaoge Nian, Yijing Cen.

Writing-review & editing: Jiayun Li, Paul Holford, George Andrew Charles Beattie, Xiaoge Nian, Yijing Cen.

## Competing interests

The authors declare no conflict of interest.

## Data Availability Statement

The published article includes all data generated or analyzed during this study. The full sequences were submitted to NCBI, accession number: MG550150.1 for *DcAKH*, OR259432 for *DcAKHR*, OP251123 for *DcMet*, XM_026820026.1 for *DcKr-h1*, XM_008488883.3 for *DcVg-1-like*, XM_026832896.1 for *DcVg-A1-like* and OP251122 for *DcVgR*.

## Supplemental Information

**Figure S1.**
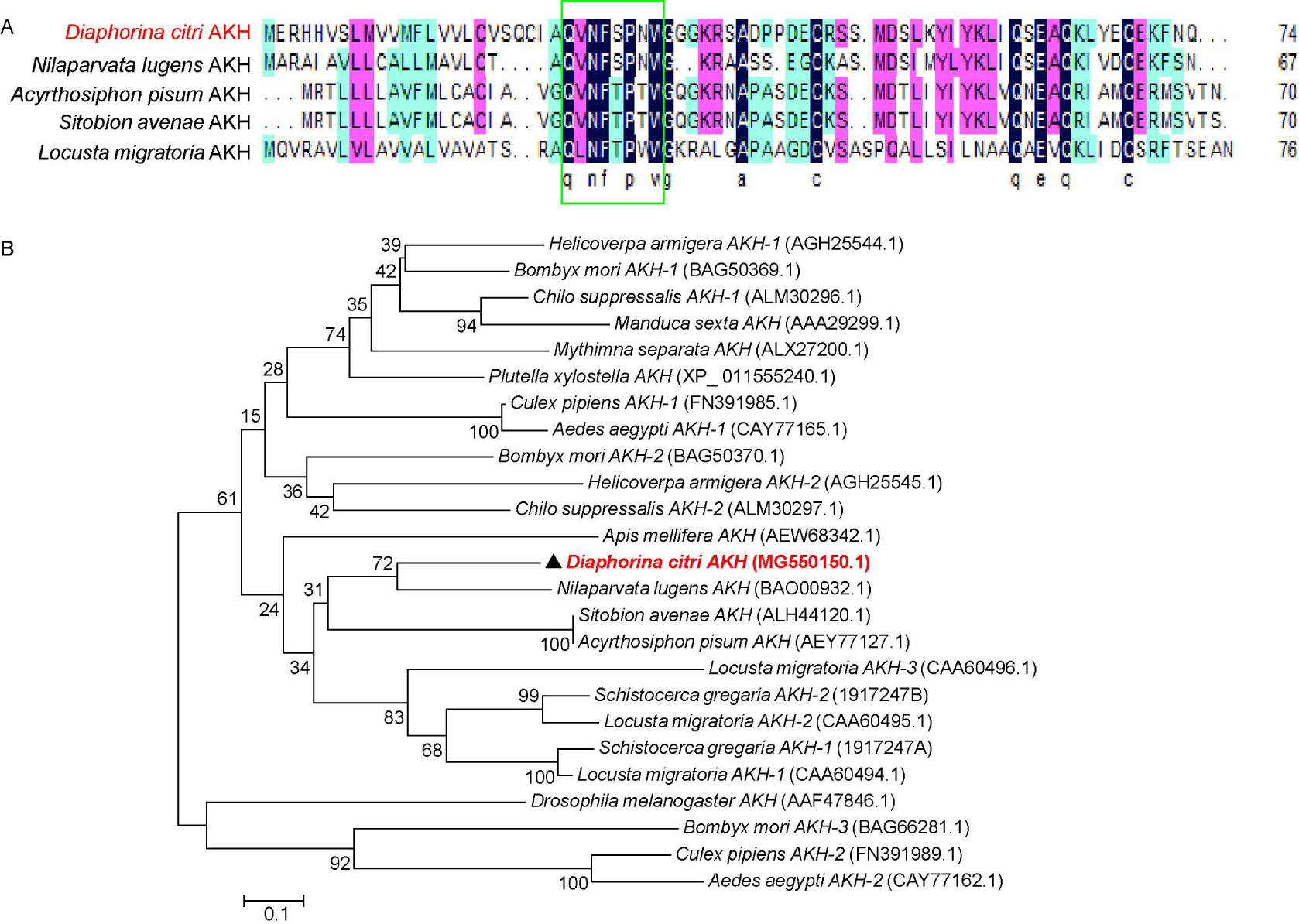
Sequence characterization of AKH from different insect species (data are related to Figure 2). (A) Alignment of the amino acid sequences of the *Dc*AKH transmembrane domain with homologs from other insect species. The sequences in the green box represent the mature peptides (QVNFSPNW). (B) Phylogenetic analysis of *Dc*AKH and its homologs in other insect species. The phylogenetic tree was constructed using neighbor-joining with 1000 bootstrap replicates; values > 50% are shown on the tree. The scale bar indicates the number of amino acid substitutions per site.

**Figure S2.**
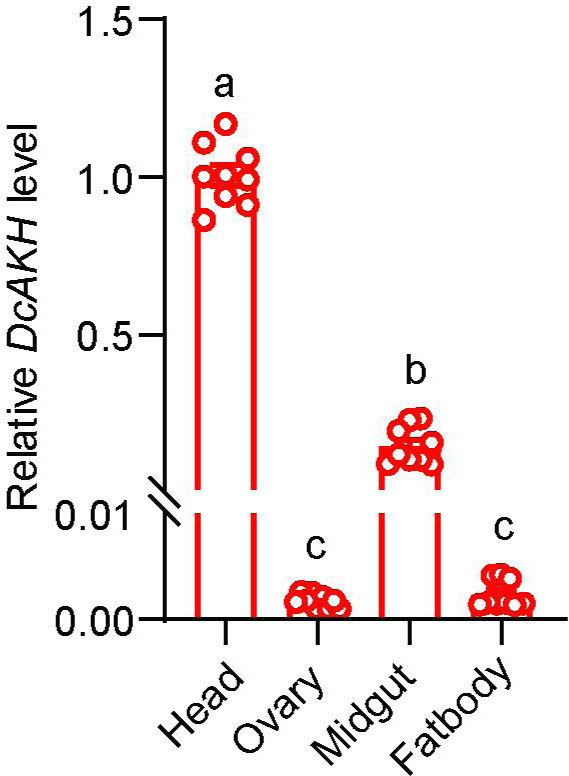
Tissue expression of *DcAKH* in *C*Las-positive adult females 9 days after emergence in the head, ovary, fat body, and midgut (data are related to Figure 2).

**Figure S3.**
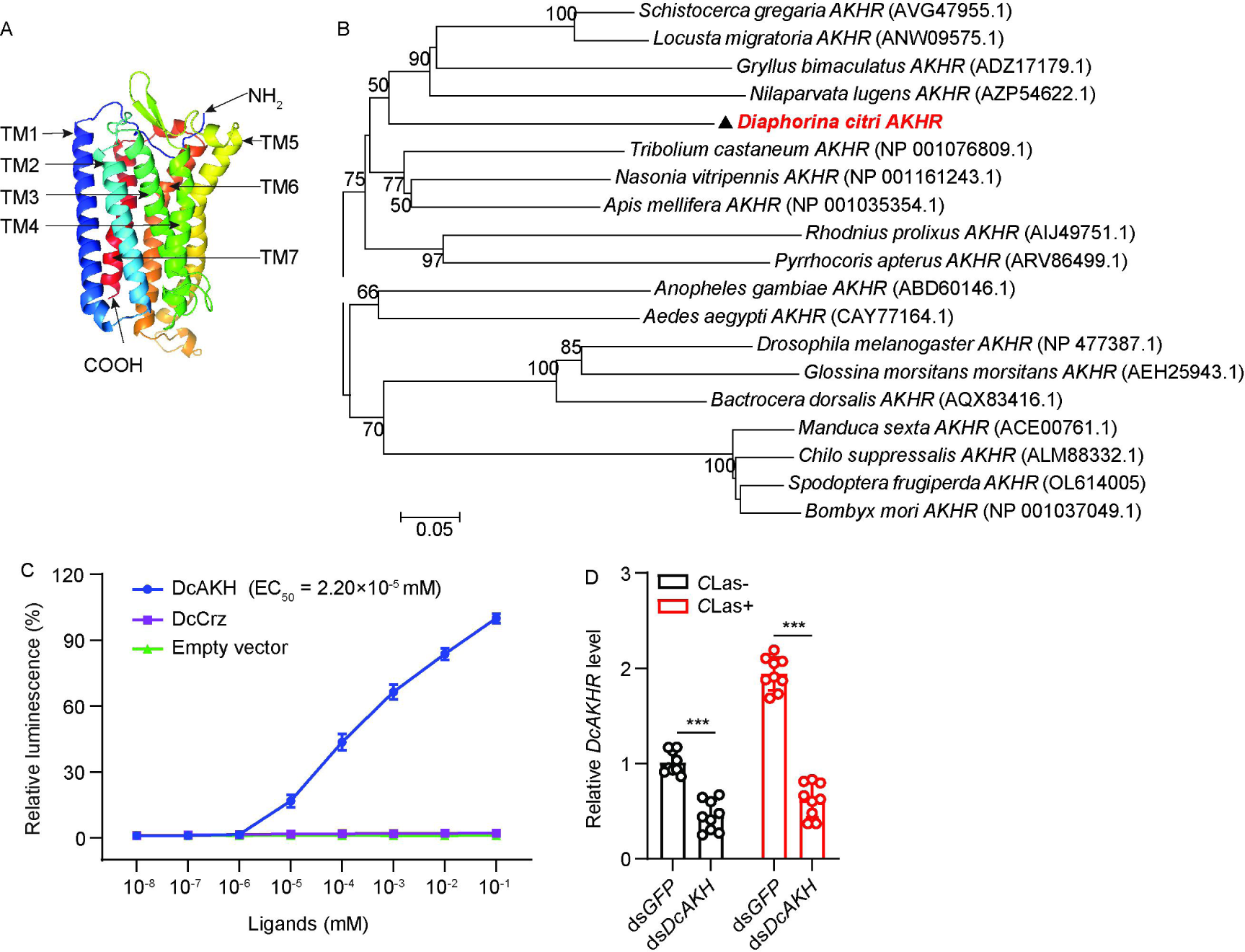
*In vivo* and *in vitro* studies validating the DcAKH-DcAKHR interaction (data are related to Figure 3). The predicted tertiary structure of the *DcAKHR* protein. The seven conserved transmembrane ion channels are shown as TM1-TM7. (B) Phylogenetic analysis the protein sequences of AKHR. The phylogenetic tree was constructed using neighbor-joining with 1000 bootstrap replicates; values > 50% are shown on the tree. The scale bar indicates the number of amino acid substitutions per site. (C) Concentration-response curves for the effect of Ca^2+^ on *DcAKHR*-expression in CHO cells. (D) The expression levels of *DcAKHR* in *C*Las-negative and *C*Las-positive females treated with ds*DcAKH* for 48 h.

**Figure S4.**
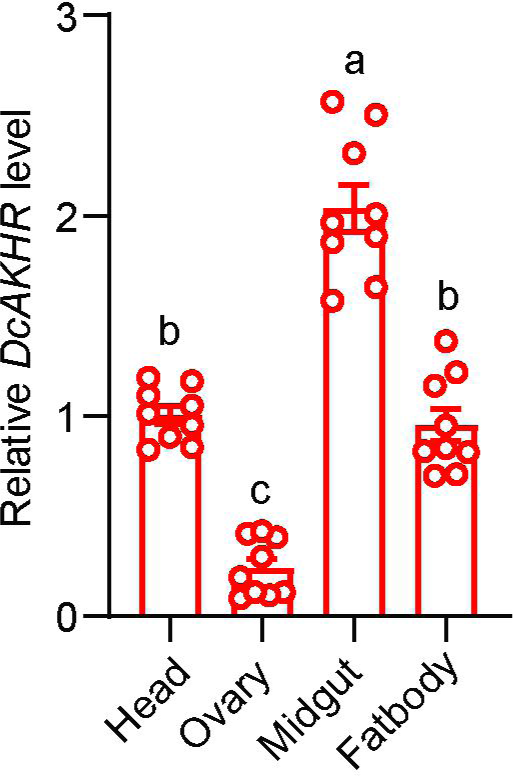
Tissue expression of *DcAKHR* in *C*Las-positive female adults 9 DAE in the head, ovary, fat body, and midgut (the data are related to Figure 3).

**Figure S5.**
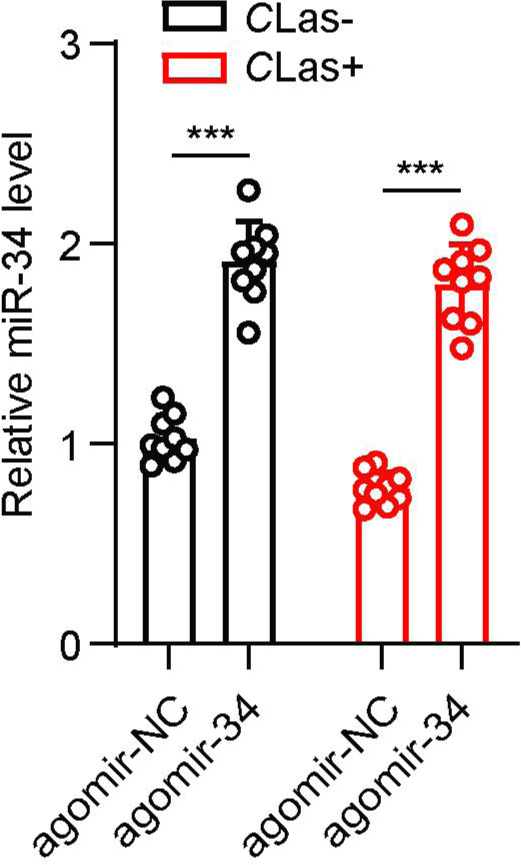
The expression levels of miR-34 in *C*Las-negative and *C*Las-positive females treated with agomir-34 for 48 h (the data are related to Figure 4).

**Figure S6.**
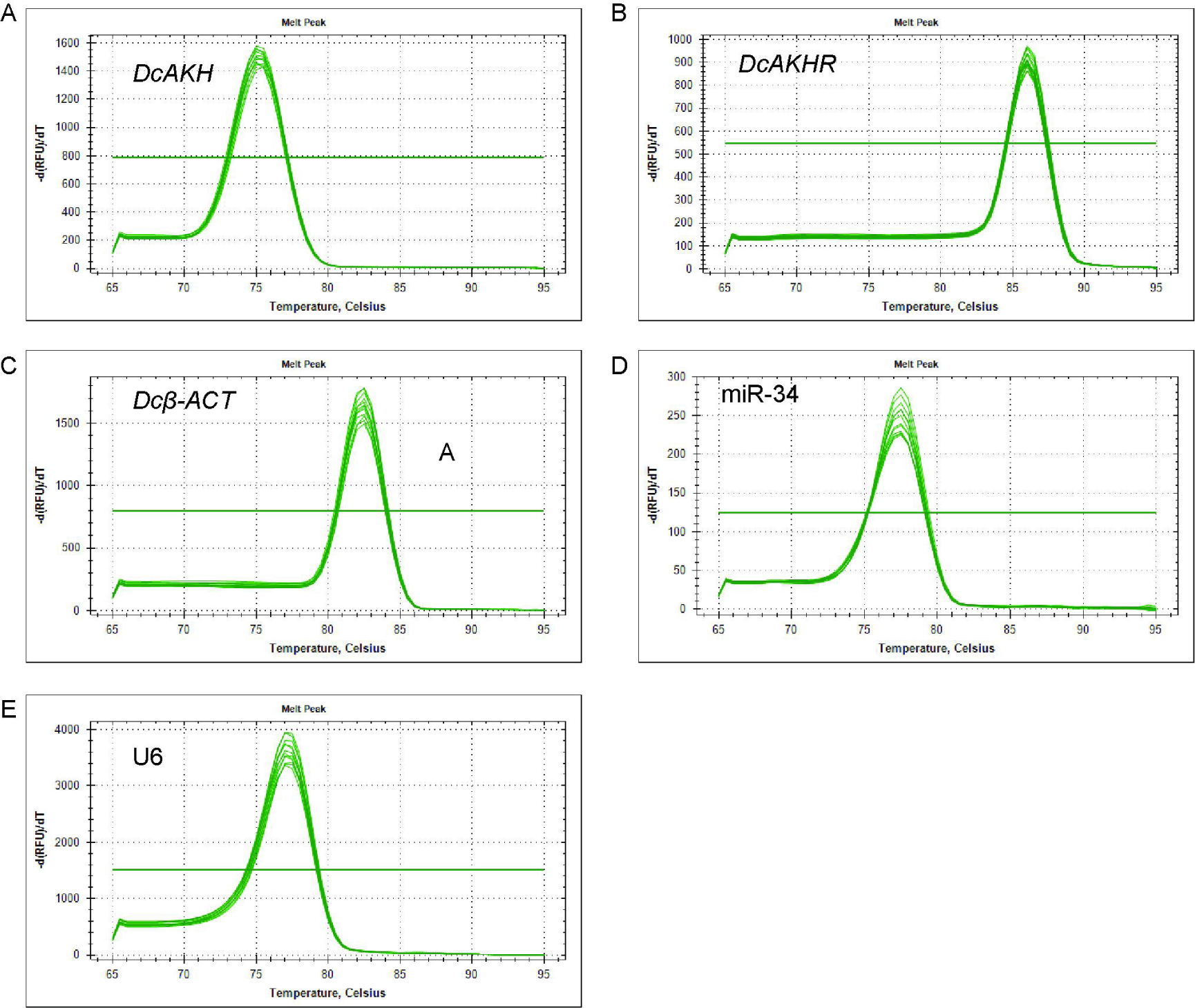
(A-E) Melting curve for qRT-PCR primers of *DcAKH*, *DcAKHR*, *Dcβ-ACT,* miR-34 and U6.

**Table S1.**
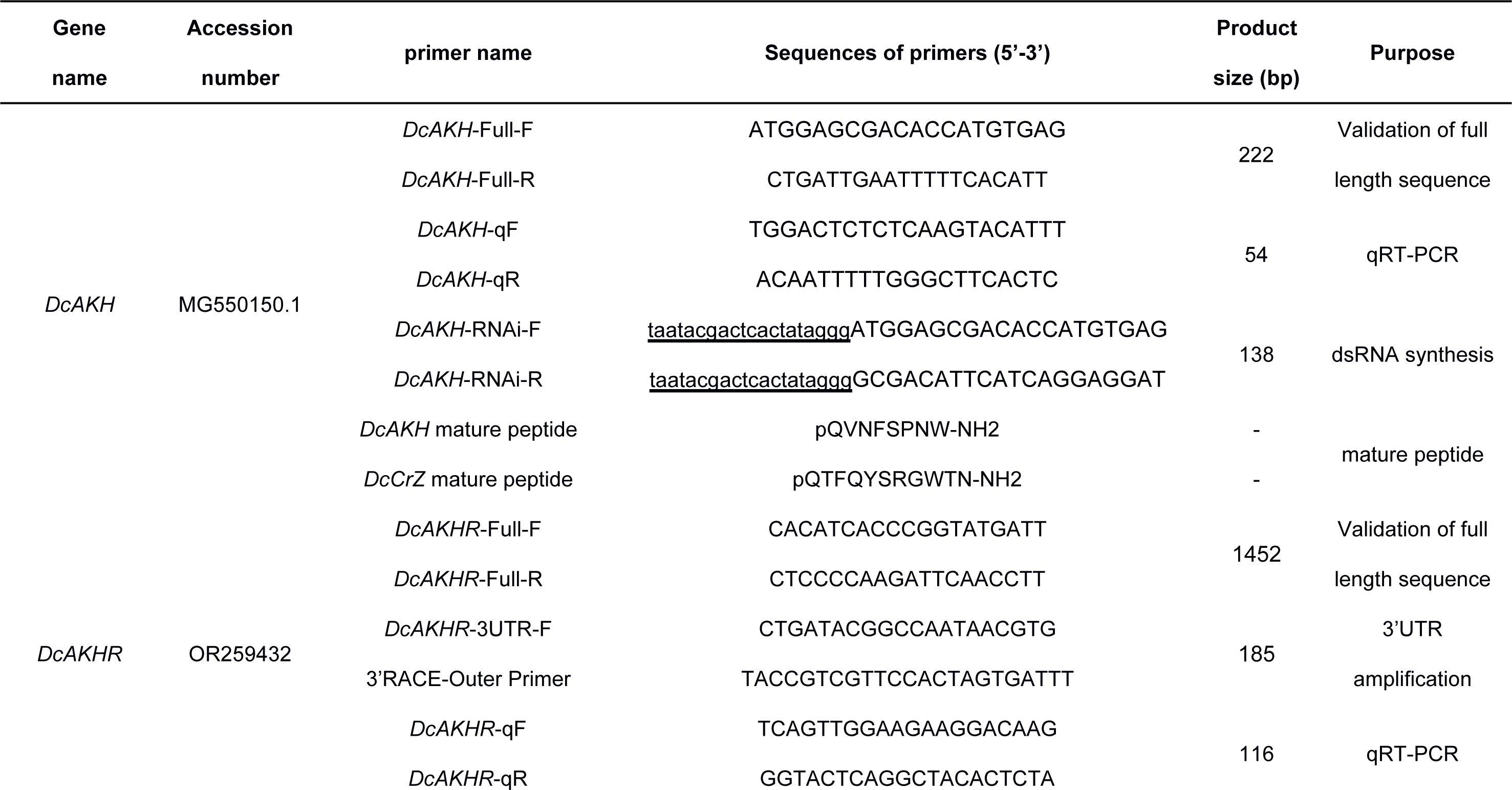
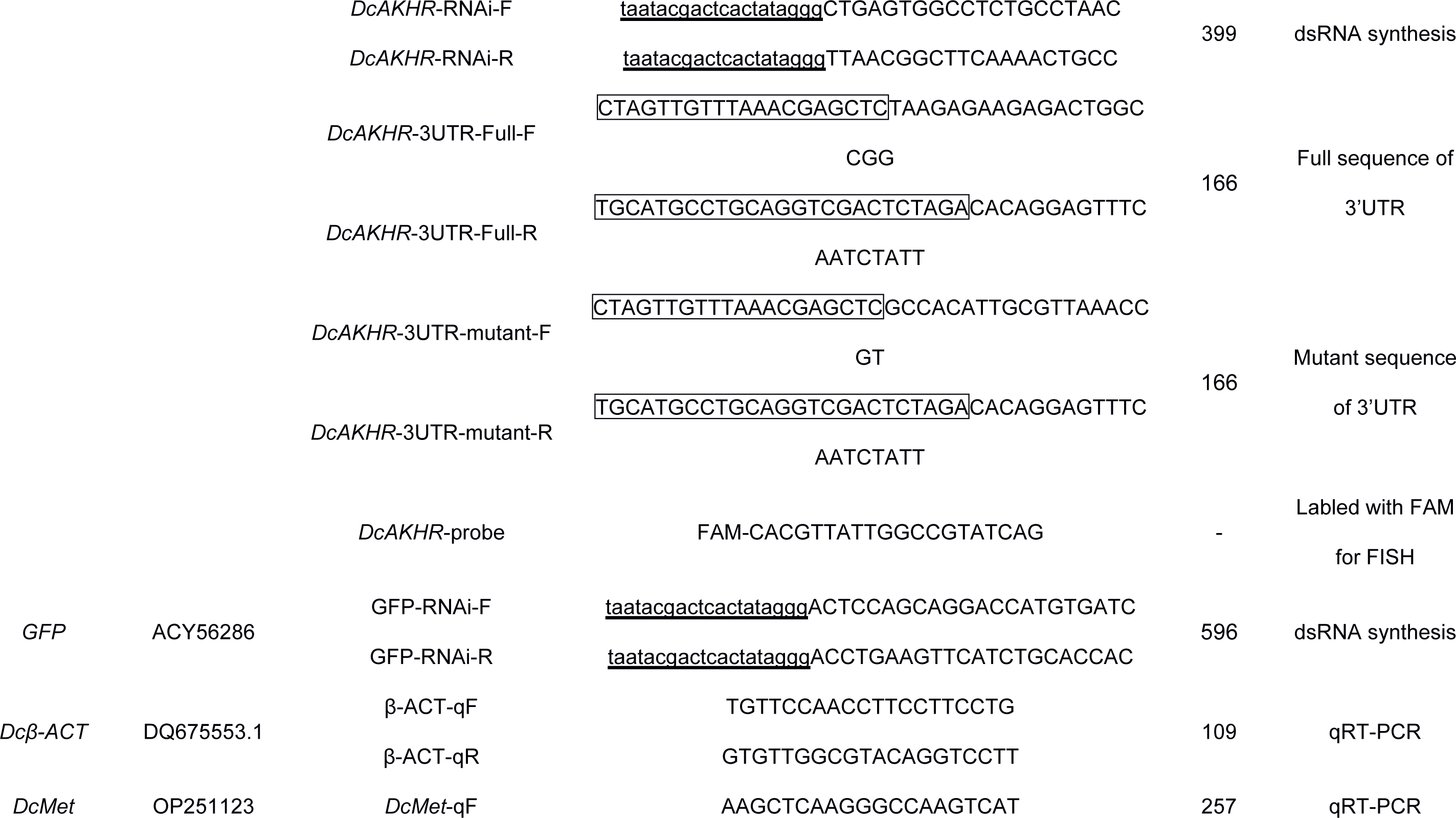
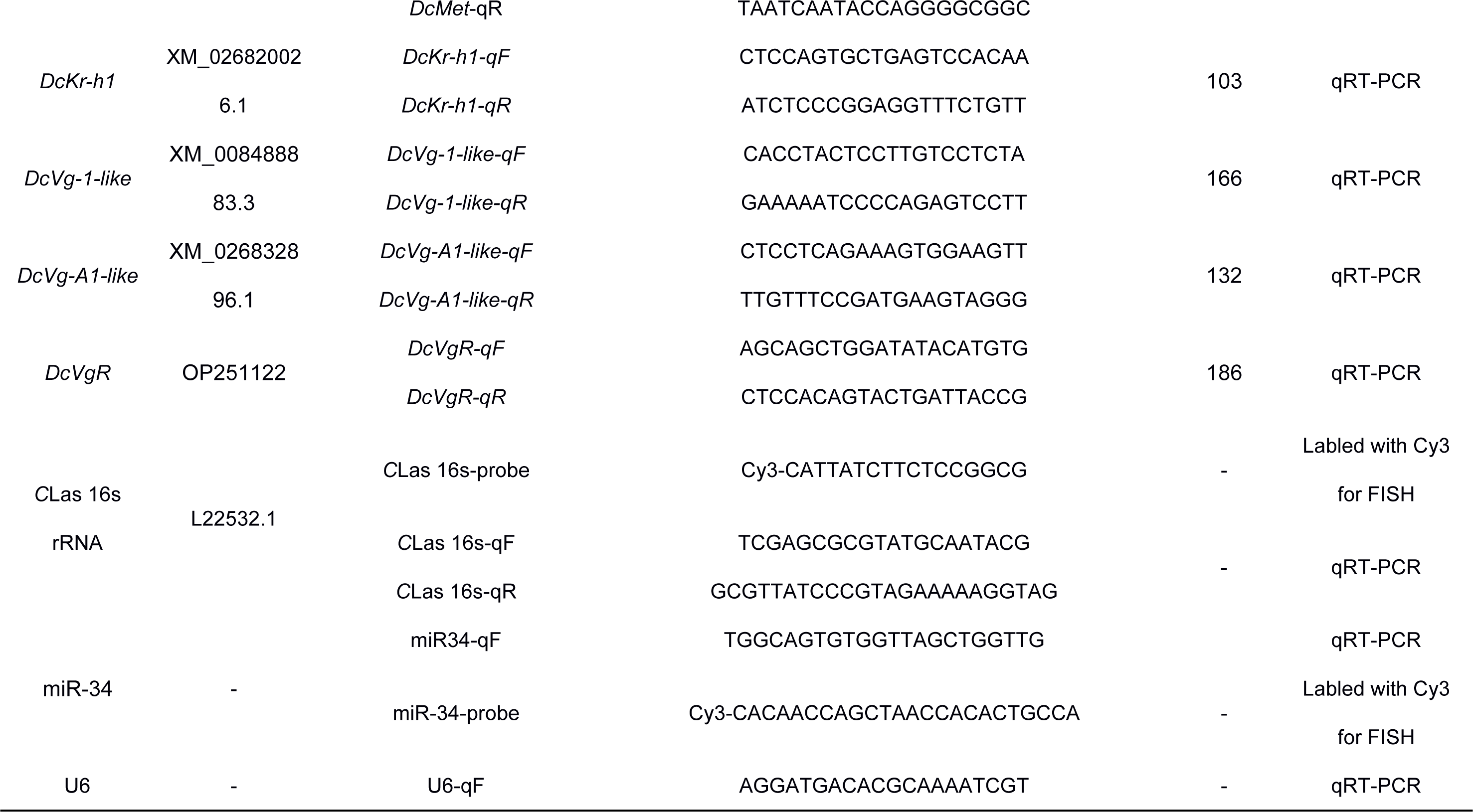
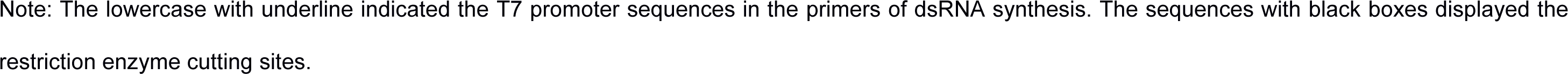
Primer list used for 3’RACE, qRT-PCR, and RNAi analysis. Related to Method details.

## Notes

### Competing Interest Statement

The authors have declared no competing interest.

### Summary of Updates

Title revised; figure 1D-E revised; introduction and discussion updated; the digit of the scale bars for Nile red staining were added in the Figure 1C, 2E, 3E, 5C; figure S6 was added.

